# HLAfreq: Download and combine HLA allele frequency data

**DOI:** 10.1101/2023.09.15.557761

**Authors:** David A. Wells, Michael McAuley

## Abstract

The Allele Frequency Net Database contains publicly available data on human immune gene frequencies for different populations from a variety of sources. We introduce a python package `HLAfreq` to download, combine, and analyse this data. In particular, `HLAfreq` can combine multiple datasets and/or prior knowledge to improve allele frequency estimates and produce credible intervals. We demonstrate these features using real world data to visualise allele frequencies and compare populations. `HLAfreq` is hosted at www.github.com/Vaccitech/HLAfreq/ with documentation and detailed examples.

## 1 Introduction

Human leukocyte antigen (HLA) genes encode cell-surface proteins which play an important role in immunity. These HLA proteins are responsible for presenting pathogen-derived peptides to T cells, an essential step in T cell activation. HLA genes are highly polymorphic and so the population frequency of HLA alleles is often considered when designing vaccines [5]. Specific HLA alleles have been linked to autoimmune disease [15] and associated with adverse drug reactions [2]. Further, the success of solid organ and stem cell transplants is related to HLA matching between donor and recipient [10, 3].

The Allele Frequency Net Database (www.allelefrequencies.net) is a publicly available repository for human immune gene frequency data from across the world [4]. It is a key resource for medical and scientific research related to HLA allele frequencies. However, difficulties downloading and combining data from multiple studies make it hard for researchers to study larger regions or even single countries where the data is split across many sources.

Here we present `HLAfreq`: a python package which can be used to download, combine and analyse datasets from the Allele Frequency Net Database. This tool should be easy to use for anybody with basic python skills, and considerably faster and more powerful than the manual alternative. To get started see the guide and examples at github.com/Vaccitech/HLAfreq. `HLAfreq` provides functions to identify incomplete studies, harmonise allele resolution, calculate population coverage, and estimate allele frequencies using a Bayesian framework. Allele frequency plots can be generated to identify anomalous datasets and interesting diversity in a set of populations. Below we describe the framework used to combine population datasets in `HLAfreq`, illustrate the basic use case, and demonstrate how `HLAfreq` can be used to study global differences in HLA allele frequencies.

## 2 Methods

### 2.1 Statistical methods

`HLAfreq` uses a Bayesian framework to estimate allele frequency statistics from combined datasets for a specific population. The user can select from two possible statistical models. The simpler ‘default model’ gives point estimates for allele frequencies. The more sophisticated ‘compound model’ gives both point estimates and credible intervals. Credible intervals can be thought of as ‘the Bayesian equivalent of confidence intervals’ although they have different mathematical interpretations; see [7, Chapter 7.7] for details.

#### 2.1.1 Default model

Let *p*_*k*_ be the frequency of the *k*-th allele of a particular gene in a given population (e.g. a country). The default model assumes that the observations from all datasets for the population are drawn independently and that the probability of being the *k*-th allele is *p*_*k*_. In other words each observation is drawn from a categorical distribution with parameters (*p*_1_, …, *p*_*K*_) where *K* is the total number of alleles. The prior for (*p*_1_, …, *p*_*K*_) is taken to be a Dirichlet distribution with parameters *α*_1_, …, *α*_*K*_ which are chosen by the user. The Dirichlet distribution is a generalisation of the Beta distribution to higher dimensions; its properties are described in [11, Section 4.6.3].

The Dirichlet distribution is conjugate to the categorical distribution, meaning that the posterior distribution for the default model is also Dirichlet. More precisely, if the combined datasets contain *x*_*k*_ observations of the *k*-th allele (for *k* = 1, …, *K*) then the posterior distribution is Dirichlet with parameters *α*_1_ + *x*_1_, …, *α*_*K*_ + *x*_*K*_. The posterior mean for the frequency of allele *j* is then given by

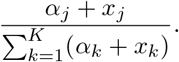

By default, `HLAfreq` takes the prior parameters to be *α*_1_ = … = *α*_*K*_ = 1. This results in a uniform prior on (*p*_1_, …, *p*_*K*_) subject to the constraints that *p*_1_, …, *p*_*K*_ *≥* 0 and *p*_1_ + … + *p*_*K*_ = 1. The user can specify alternative values for *α*_1_, …, *α*_*K*_. These parameters may be interpreted as a ‘pseudocount’ in the sense that choosing the prior *α*_1_, …, *α*_*K*_ is equivalent to taking a uniform prior and then observing a dataset with *α*_*k*_ *−*1 observations of the *k*-th allele for *k* = 1, …, *K*. (Intuitively the uniform prior corresponds to one observation of each allele). This can be used as a heuristic for choosing prior parameters based on external information.

The default model could in principle be used to estimate credible intervals for the frequencies *p*_1_, …, *p*_*K*_ however this option is not provided by `HLAfreq` because in practice such intervals are frequently unrealistically narrow. This is because the default model does not account for variance between studies. In the next subsection we outline how we can account for this variation and obtain accurate credible intervals. The current model is chosen as the default because it is simpler and we expect its point estimates to be sufficient for the majority of use cases.

#### 2.1.2 Compound model

The default model assumes that all observations are independent and sampled without bias; however, observations within a single study are more likely to be similar e.g. they may be sampled at the same time or place. To account for this, `HLAfreq` provides a ‘compound model’ which accounts for the grouping of observations within studies and allows the allele frequencies of studies to differ from each other. The additional uncertainty of the model results in wider credible intervals which better reflect the true state of knowledge regarding population frequencies.

The compound model makes the following assumptions. As before, *p*_*k*_ denotes the frequency of the *k*-th allele in the population and the prior distribution for *p*_1_, …, *p*_*K*_ is Dirichlet with parameters *α*_1_, …, *α*_*K*_. A concentration parameter *γ≥*0 is given with a standard log-normal prior distribution.

For the *j*-th data source, a vector 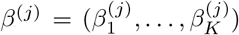 is sampled independently from a Dirichlet distribution with parameters *γp*_1_, …, *γp*_*K*_. Observations from the *j*-th data source are then sampled from a categorical distribution with parameters 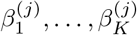. (Equivalently, the *j*-th data source as a whole is sampled from a multinomial distribution.)

Idiosyncratic sampling biases are captured by the different values of *β*^(*j*)^, which result in different probabilities of sampling particular alleles for each data source. If *γ* is large, then *β*^(*j*)^ is likely to concentrate around (*p*_1_, …, *p*_*K*_) which means that different studies tend to have similar allele frequencies.

The posterior distributions of *p*_1_, …, *p*_*K*_ and *γ* do not have a closed form and so are estimated numerically using `PyMC` [13]. The `HLAfreq` function `AFhdi` outputs posterior means and credible intervals for allele frequencies.

### 2.2 Software details

`HLAfreq` provides a simple python interface to the Allele Frequency Net Database’s Classical HLA allele frequency search and all its filters [4]. The data is downloaded by “web scraping” the Allele Frequency Net Database as described in their “automated access” section. The search results are downloaded as a pandas dataframe [9] which can be saved and opened like any other spreadsheet.

Any data filter on Allele Frequency Net Database is available through `HLAfreq`, so you can download just datasets from specific countries, regions, ethnicities, study types, and data sources. All filters and how to use them are described in the `HLAfreq` documentation under [makeURL](https://github.com/Vaccitech/HLAfreq/blob/main/docs/HLAfreq.md#function-makeurl-hlafreqhlafreqmakeurl). Class I and class II alleles can be downloaded by specifying the locus name. The Allele Frequency Net Database characterises each dataset as gold, silver, or bronze standard. You can specify the minimum dataset quality to download by passing `“g”`, `“s”`, or `“a”` (all) to the `standard` argument of `makeURL()`.

The function `combineAF()` is used to combine multiple datasets and estimate the allele frequencies according to the default model described above. So that larger studies contribute more to the combined allele frequency estimate, each dataset is weighted by twice the sample size by default (because each individual is diploid). Alternatively any supplied weighting can be used; for example, population size can be used for a multi-country estimate. If credible intervals are required for a multi-country estimate, then the compound model should be used for each country before weighting by population size. This is demonstrated in the [multi-country example](https://github.com/Vaccitech/HLAfreq/blob/main/examples/multi_country.ipynb). `combineAF()` performs several automated checks on the dataset which can also be performed using provided helper functions. Studies are flagged as incomplete if the total allele frequency is outside a specified range (0.95-1.1 by default). Unmeasured alleles are added with a frequency of zero to ensure all populations report the same set of alleles before combining. HLA alleles can be reported to different resolutions but only alleles of the same resolution can be combined; therefore, allele resolution is automatically checked when combining datasets. By default `combineAF()` uses a flat prior of 1 observation for each allele but this can be set manually.

The allele frequency estimates and credible intervals for the compound model can be obtained with `AFhdi()` imported from the submodule `HLAfreq.HLAfreq_pymc`.

Once allele frequencies have been combined they can easily be plotted with existing plotting tools such as matplotlib’s `bar()` [6]; however, we also provide `plotAF()` to view the combined allele frequency estimates, credible intervals, and the original values of each dataset.

## 3 Results

In this section we illustrate the use of `HLAfreq` through several examples.

The following code downloads and combines all gold standard datasets for Mongolia at HLA locus DQB1 and plots the population allele frequencies from both the default and compound model, and plots the credible interval from the compound model (see Figure 1).

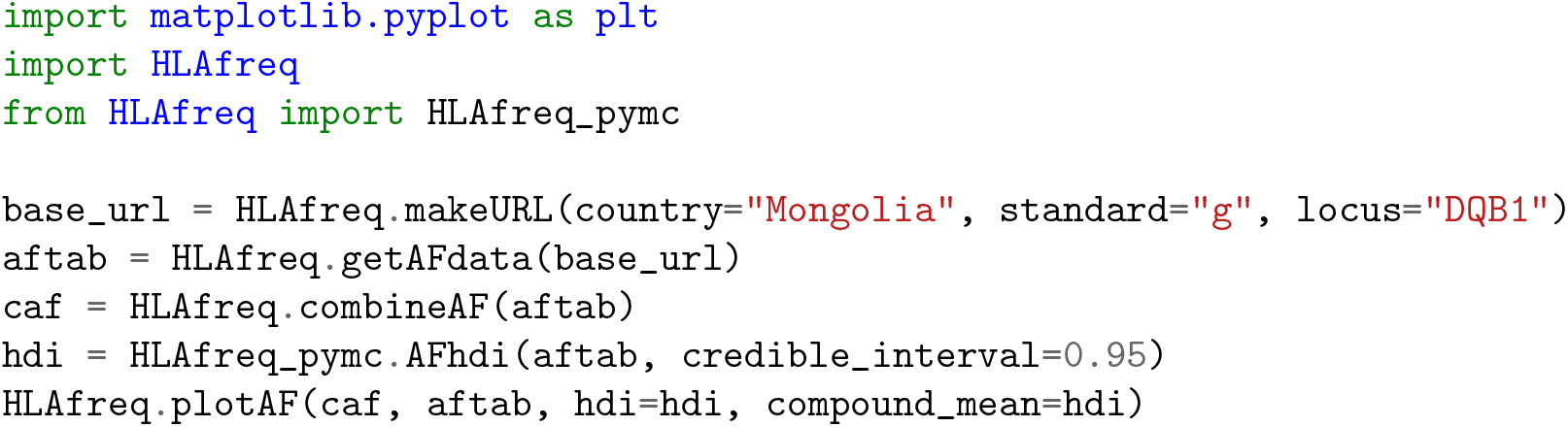

**Figure 1.**
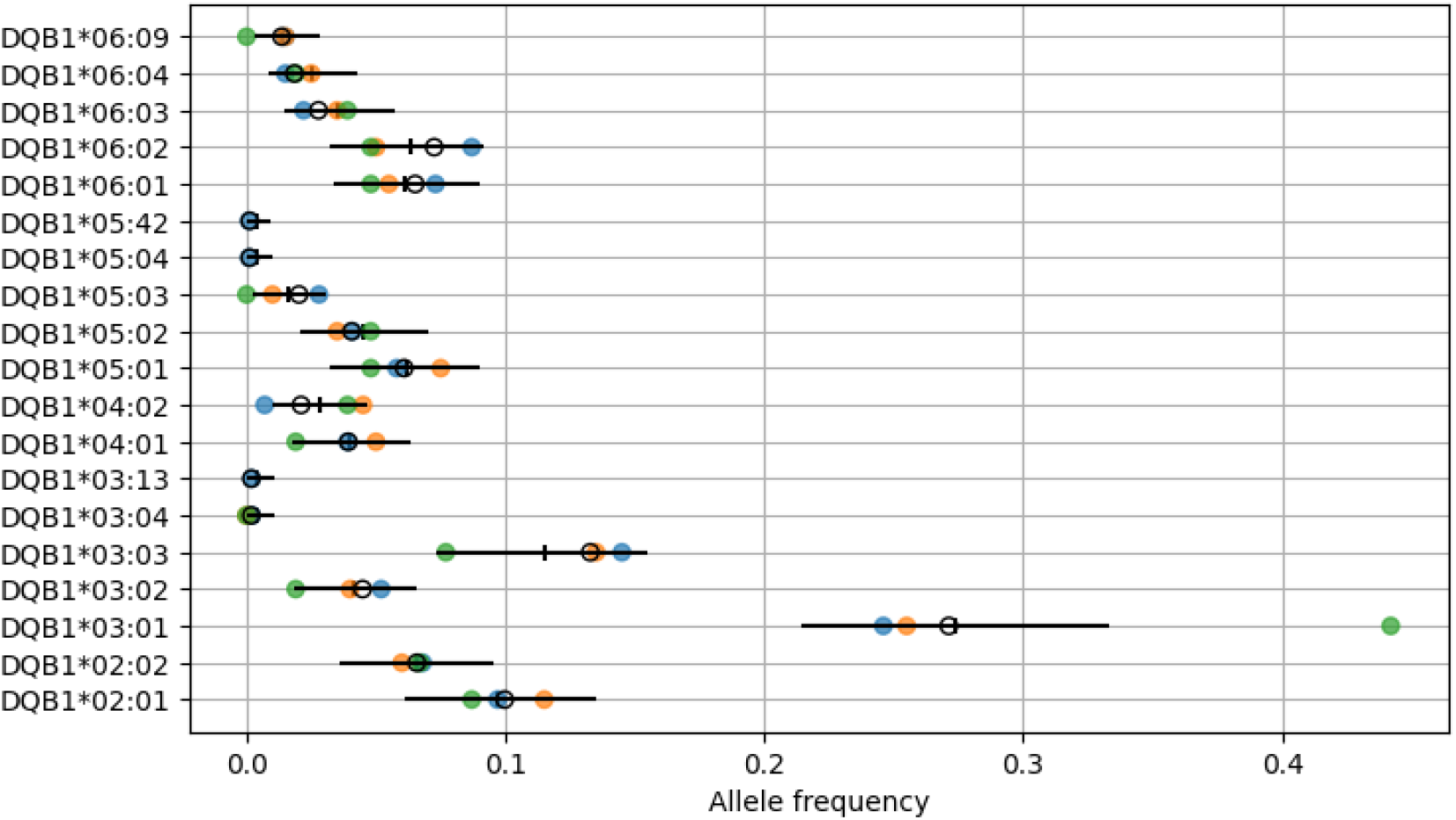
Allele frequencies for gold standard datasets of the Mongolian population at HLA locus DQB1. Default estimated population frequencies are shown in open black circles and observed frequencies from individual studies are shown in closed circles of different colours. Estimated allele frequencies and credible intervals from the compound model are shown as vertical and horizontal bars respectively.

The allele frequencies of individual studies are included in the plot to help identify heterogeneity between datasets (e.g. DQB1*03:01 in Figure 1).

After combining datasets, the population coverage of any allele or set of alleles can be estimated using `population_coverage()`. Population coverage is calculated under the assumptions of Hardy-Weinberg equilibrium as *p*^2^ + 2*p*(1*−p*) where *p* is the cumulative frequency of a set of alleles. The following code computes and plots the cumulative population coverage of the most frequent alleles for the Mongolian dataset used above (see Figure 2).

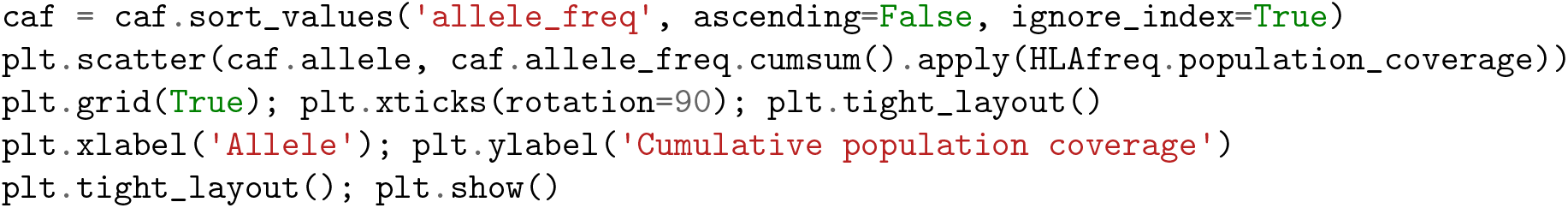

**Figure 2.**
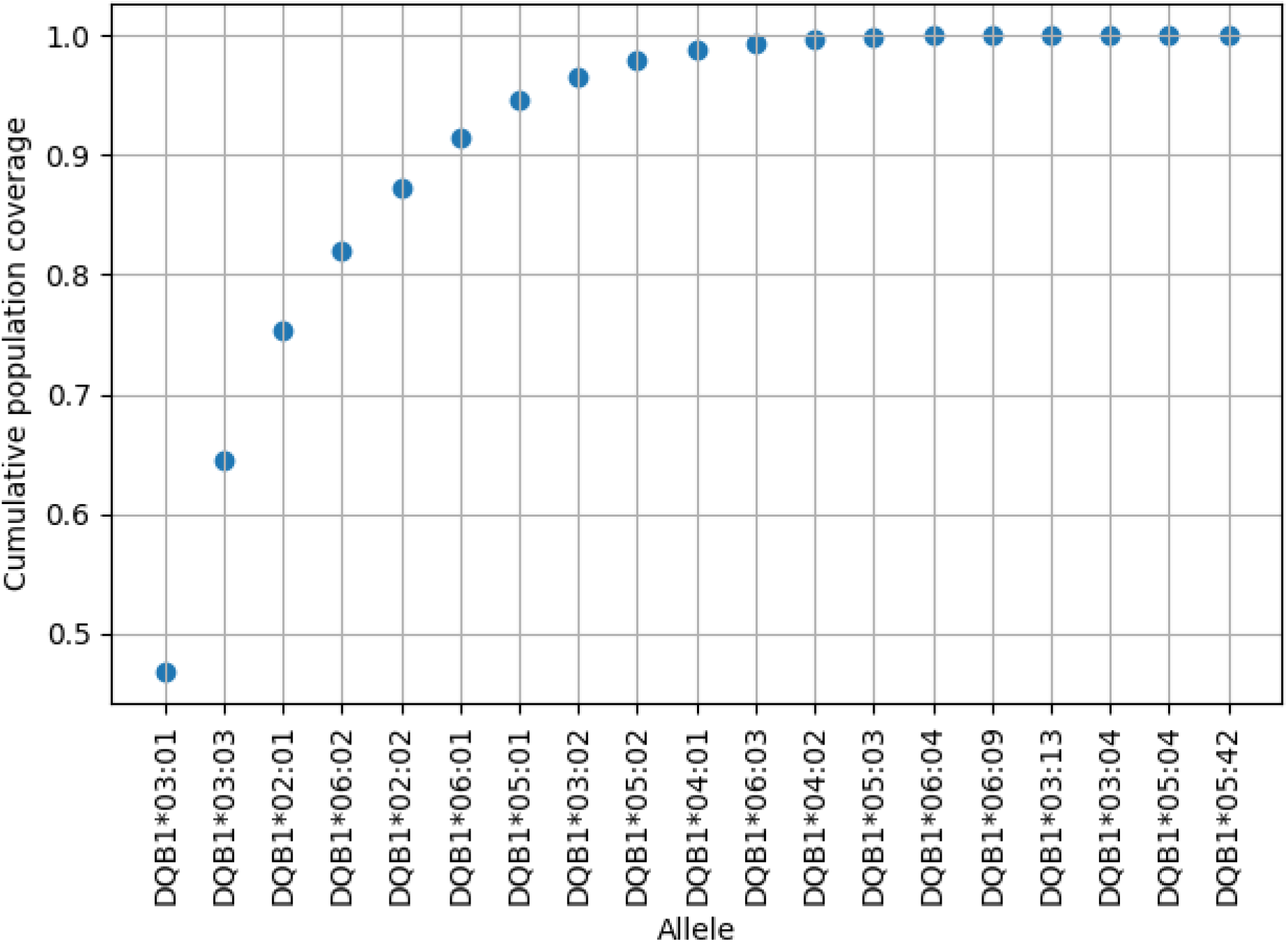
Estimated cumulative population coverage of the most frequent alleles at HLA locus DQB1 for Mongolia.

This function can be used to see how well a panel of HLA alleles covers a population or to select a panel that provides good coverage for a specific population. For instance the immune epitope database (IEDB) provides a reference panel of 27 HLA class I alleles which should cover at least 97% of the global population [16, 8]. Using `HLAfreq` we calculated the population coverage of this panel for each country and found that coverage in some countries is substantially lower (see Figure 3). Furthermore, better coverage is possible with fewer alleles if they are selected based on country specific frequency data (see Figure 4). A smaller panel can greatly reduce computation time when predicting epitopes because predictions are made per allele. This is particularly useful in vaccine design when searching for epitope rich regions or assessing the coverage of a proposed vaccine in a specified population. When designing a panel of HLA alleles to represent a population it is worth considering multiple genes and supertypes within genes rather than solely the most common alleles [14].

**Figure 3.**
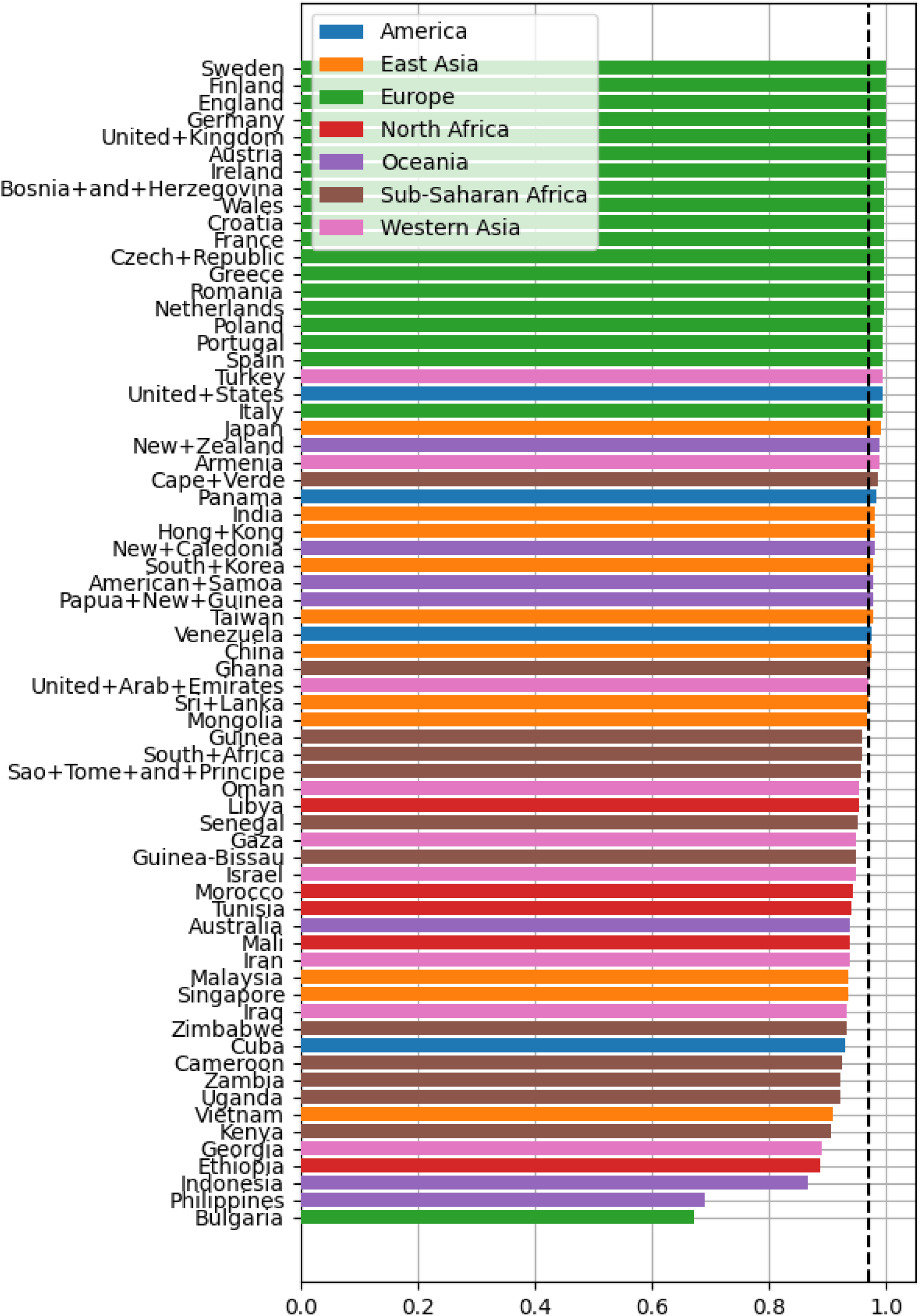
Population coverage of IEDB 27 allele panel assuming no linkage disequilibrium between HLA-A and HLA-B. Dashed line shows expected global population coverage (97%).

**Figure 4.**
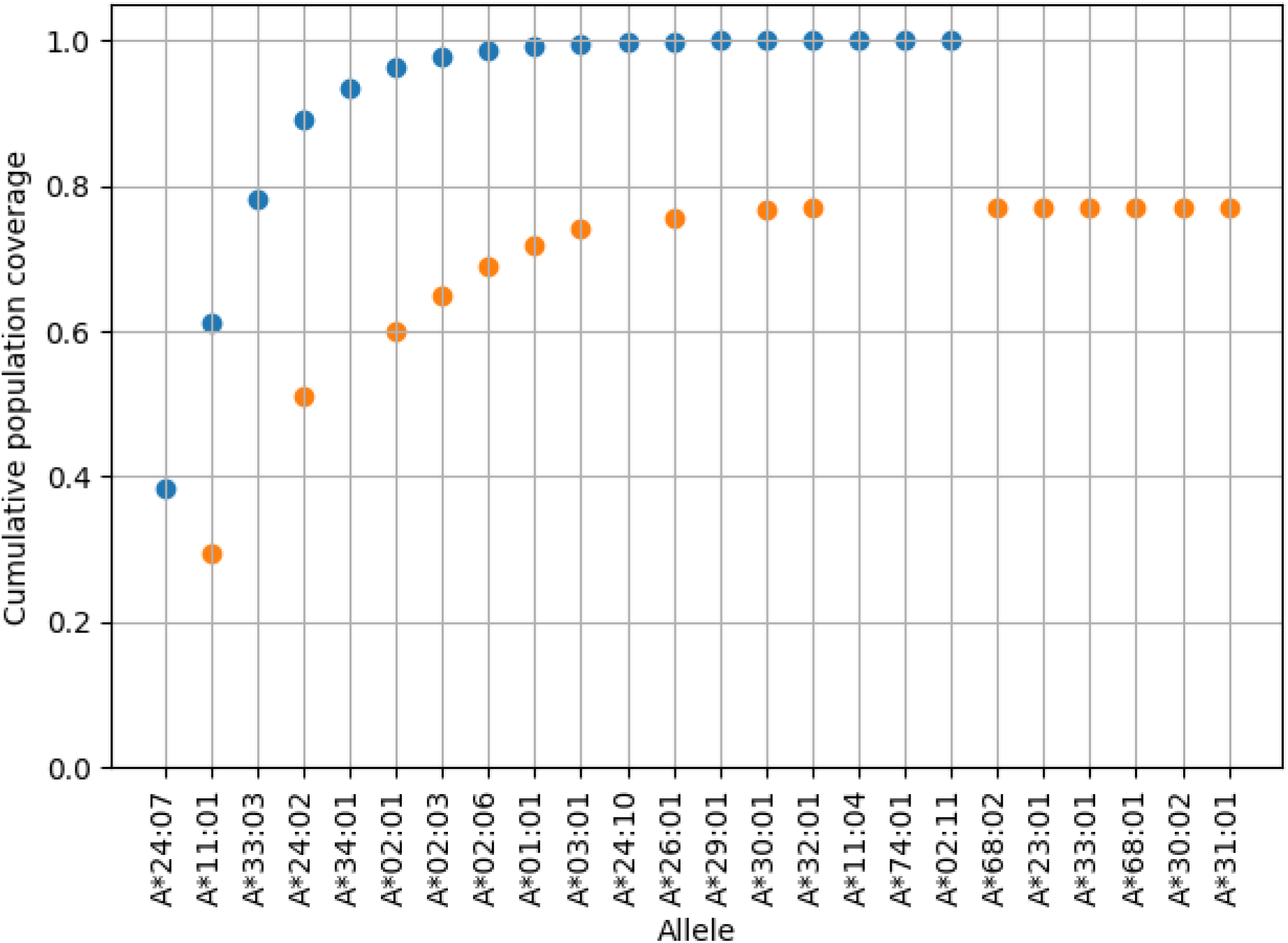
Dark blue points show cumulative frequency of all alleles observed in Indonesia. Light orange points show cumulative population coverage of only HLA A alleles in the IEDB reference panel. Alleles that are not observed in Indonesia are not plotted in blue and alleles that are not part of the IEDB reference panel are not plotted in orange.

`HLAfreq` can also facilitate cross-country comparisons. To demonstrate this, we estimated the HLA-A allele frequencies for each country by downloading and combining all available gold standard datasets. We then performed principal component analysis using scikit-learn to project allele frequencies onto two dimensions [12]. This requires each country to report the same set of allele frequencies, and so unreported alleles were added as zero using `unmeasured_alleles()`. The plotted projection (see Figure 5) shows clear grouping by geographic region, indicating similarities in HLA-A frequencies between countries in the same region.

**Figure 5.**
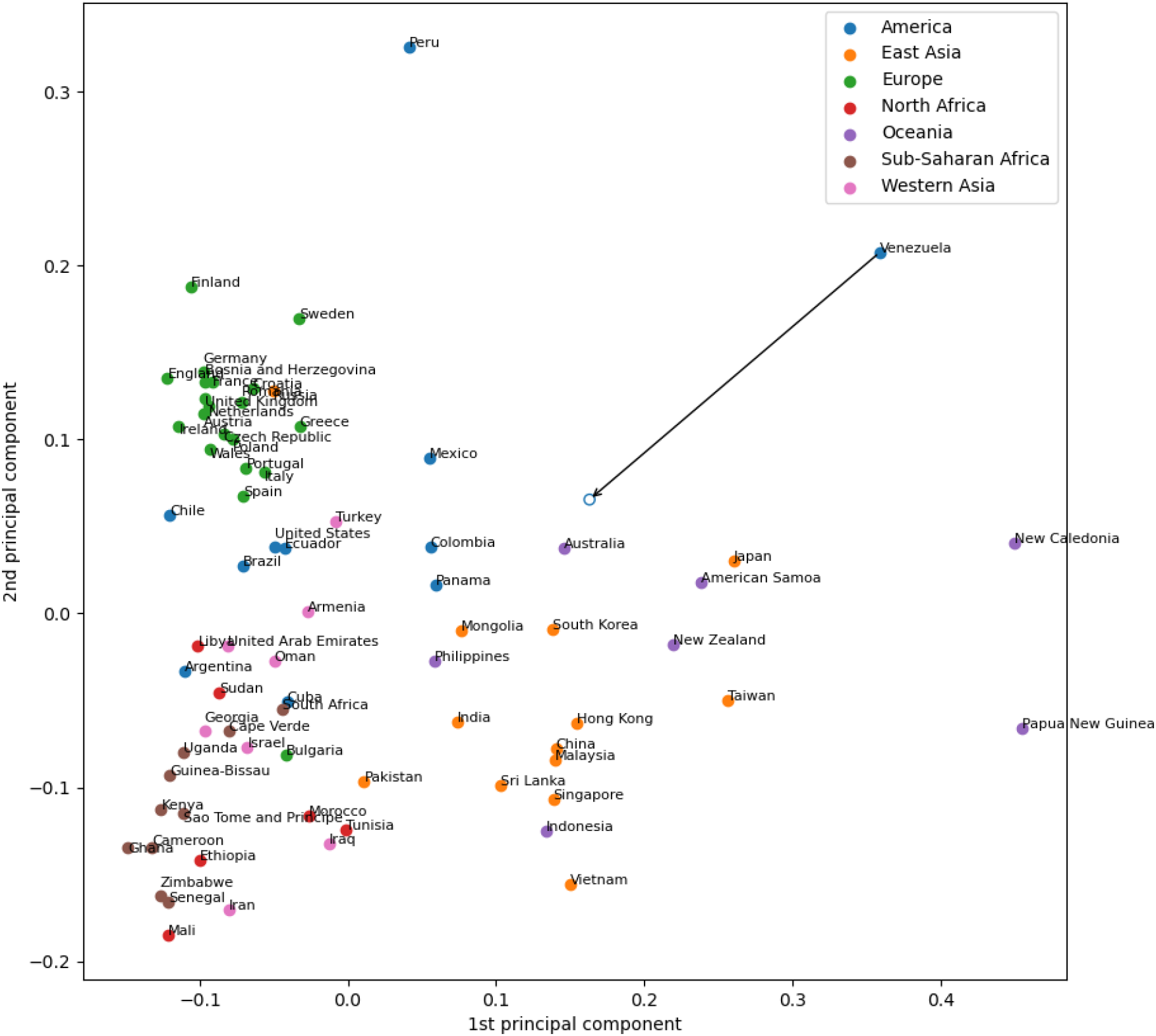
PCA of estimated allele frequencies across countries. The arrow illustrates the effect of an informative prior on Venezuela.

On the basis of this, it may be possible to improve frequency estimates for undersampled countries by using an informative prior based on neighbouring countries. Venezuela is the country with smallest sample size (55 observations) in our combined dataset. The effect of using neighbouring Colombia (1515 observations) to form a prior is illustrated by the arrow in Figure 5. For this prior, the sample size of Colombia was divided by 30 so that approximately equal information was provided by the Colombian and Venezuelan datasets. The new Venezuelan estimates appear more similar to those of Colombia and other American countries.

## 4 Discussion

We present `HLAfreq` as a fast and easy-to-use tool for downloading and combining the HLA frequency data stored in the Allele Frequency Net Database. This tool streamlines the process of working with HLA frequency data across large regions and multiple populations. `HLAfreq` can be used to study regional and global differences in HLA frequencies, and also paired with other datasets to study the impact of HLA frequencies on other immune conditions. Once HLA data has been combined at the appropriate level, e.g. national, it is straightforward to pair that data with any immune data of interest such as national rates of coeliac disease, or frequencies of particular viral variants. `HLAfreq` can be used in several types of analysis, not only when testing associations with specific alleles, but when statistically controlling for the frequency of alleles while investigating other variables, or even the effect of allelic diversity itself.

`HLAfreq` can incorporate prior knowledge to improve frequency estimates in undersampled populations. Prior weights can be specified manually as *α*_1_, …, *α*_*K*_ with the interpretation that allele *k* has previously been observed *α*_*k*_*−*1 times. Often the prior information will come from studies in neighbouring populations, in these cases it is easier to use data from those populations to generate a prior.

We have demonstrated how a prior for Venezuela can be based on neighbouring Colombia. For simplicity we used a single country but multiple can be used to generate a prior. When generating a prior from a population it is important to appropriately downweight the prior dataset so that it does not overwhelm the study dataset. In our example we have somewhat arbitrarily chosen to downweight by a factor of 30. The choice of downweighting will be study specific and depend on how similar the prior and study populations are expected to be *a priori*. When using multiple populations to generate a prior each one can be given a different weight. We recommend verifying that study conclusions are robust to the specific prior values by rerunning the analysis with alternative, reasonable priors.

When combining datasets with small total sample size but many alleles (typically *>* 100), the compound model can give estimates that are much lower than expected. This is a known issue for estimating multinomial/categorical parameters using a Bayesian paradigm [1], fortunately it is easy to diagnose and solve. Plot allele frequencies (as in Figure 1) and if compound model estimates are shifted away from observed data and towards equal frequency, use the prior *α*_1_ =… = *α*_*K*_ = 1*/K*. This prior expresses the belief that some alleles are likely to be much more common than others (without knowing which alleles these will be). A worked example of diagnosing and fixing this issue is in the [working with priors](https://github.com/Vaccitech/HLAfreq/blob/main/examples/working_with_priors.ipynb) example. The default method is far less sensitive to this issue and so `HLAfreq` produces a warning when the default and compound models diverge.

`HLAfreq` is hosted at www.github.com/Vaccitech/HLAfreq/ with documentation and detailed examples. The code for this paper is available at www.github.com/Vaccitech/HLAfreq/tree/main/examples/paper

## Acknowledgements

MM was supported by the European Research Council (ERC) Advanced Grant QFPROBA (grant number 741487). DW is employed by Vaccitech plc.

## References

[1] C. P. de Campos and A. Benavoli. “Inference with multinomial data: Why to weaken the prior strength”. Twenty-Second International Joint Conference on Artificial Intelligence. 2011.

[2] W.-L. Fan et al. “HLA association with drug-induced adverse reactions”. Journal of immunology research 2017 (2017).

[3] D. Fürst et al. “HLA matching in unrelated stem cell transplantation up to date”. Transfusion Medicine and Hemotherapy 46.5 (2019), pp. 326–336.

[4] F. F. Gonzalez-Galarza et al. “Allele frequency net database (AFND) 2020 update: Gold-standard data classification, open access genotype data and new query tools”. Nucleic Acids Research 48.D1 (2020), pp. D783–D788.

[5] K. Gulukota and C. DeLisi. “HLA allele selection for designing peptide vaccines”. Genetic Analysis: Biomolecular Engineering 13.3 (1996), pp. 81–86.

[6] J. D. Hunter. “Matplotlib: A 2D graphics environment”. Computing in Science & Engineering 9.3 (2007), pp. 90–95.

[7] B. Lambert. “A student’s guide to Bayesian statistics”. A Student’s Guide to Bayesian Statistics (2018), pp. 1–520.

[8] S. Martini. HLA allele frequencies and reference sets with maximal population coverage. 2019.

[9] Wes McKinney. “Data Structures for Statistical Computing in Python”. Proceedings of the 9th Python in Science Conference. Ed. by Stéfan van der Walt and Jarrod Millman. 2010, pp. 56

[10] Y. Morishima et al. “The clinical significance of human leukocyte antigen (HLA) allele compatibility in patients receiving a marrow transplant from serologically HLA-A, HLA-B, and HLA-DR matched unrelated donors”. Blood, The Journal of the American Society of Hematology 99.11 (2002), pp. 4200–4206.

[11] K. P. Murphy. Probabilistic machine learning: an introduction. MIT press, 2022.

[12] F. Pedregosa et al. “Scikit-learn: Machine Learning in Python”. Journal of Machine Learning Research 12 (2011), pp. 2825–2830.

[13] J. Salvatier, T. V. Wiecki, and C. Fonnesbeck. “Probabilistic programming in Python using PyMC3”. PeerJ Computer Science 2 (2016), e55.

[14] J. Sidney et al. “HLA class I supertypes: A revised and updated classification”. BMC Immunology 9 (2008), pp. 1–15.

[15] M. Simmonds and S. Gough. “The HLA region and autoimmune disease: associations and mechanisms of action”. Current genomics 8.7 (2007), pp. 453–465.

[16] D. Weiskopf et al. “Comprehensive analysis of dengue virus-specific responses supports an HLA-linked protective role for CD8+ T cells”. Proceedings of the National Academy of Sciences of the United States of America 110.22 (2013).

